# Regulation of Hedgehog Signaling Through Arih2-Mediated Smoothened Ubiquitination and Endoplasmic Reticulum-Associated Degradation

**DOI:** 10.1101/2022.02.20.481183

**Authors:** Bo Lv, Xiao-Ou Zhang, Gregory J. Pazour

## Abstract

During Hedgehog signaling, the ciliary levels of Ptch1 and Smo are regulated by pathway activity. At the basal state, Ptch1 localizes to cilia and prevents the ciliary accumulation and activation of Smo. Upon stimulation with Hedgehog ligand, Ptch1 exits cilia and relieves the inhibition of Smo. Uninhibited Smo concentrates in cilia, becomes activated, and activates the downstream steps of the pathway. Loss of the ubiquitin E3 ligase Arih2 elevates the cellular level of Smo, causes Smo to inappropriately localize to cilia at the at the basal state, and elevates basal expression of Hedgehog responsive genes. Mice express two isoforms of Arih2 with different N-termini, but neither isoform localizes to cilia. Instead, Arih2α is found in the nucleus and Arih2β is found on the cytoplasmic face of the endoplasmic reticulum. Re-expression of endoplasmic reticulum-localized Arih2β, but not nuclear-localized Arih2α returns the cellular Smo levels back to normal and rescues the ciliary Smo accumulation phenotype. When Arih2β is missing, protein aggregates accumulate in the endoplasmic reticulum and the unfolded protein response is activated. Inhibitor studies suggest that Arih2β functions to mark excess or misfolded Smo for degradation by endoplasmic reticulum-associated degradation. When Arih2β is defective, excess Smo, possibly misfolded, is delivered to the cell surface and cilium where it interferes with pathway regulation. These findings add another level of complexity to the Hedgehog pathway.

## Introduction

The Hedgehog pathway is an evolutionarily conserved signaling cascade that functions in embryonic development and tissue homeostasis. Malfunction of the pathway causes a variety of developmental syndromes and cancers. In vertebrates, this pathway is mediated by cilia, hair-like organelles found in nearly all eukaryotic cells. In brief, at the basal state without sonic hedgehog (SHH) ligand present, the SHH receptor Ptch1 localizes to cilia and inhibits the ciliary accumulation and activation of Smo. Upon binding of the Hedgehog ligand, Ptch1 stops inhibiting Smo and Ptch1 exits cilia. Uninhibited Smo accumulates in cilia, becomes activated, and promotes the activation of the Gli transcription factors, which move to the nucleus to promote gene expression. Our previous work showed that ciliary Smo levels are regulated by ubiquitination. At the basal state, the ubiquitin E3 ligase Wwp1 localizes to cilia by binding Ptch1. This promotes the ubiquitination of Smo, which promotes Smo’s interaction with the intraflagellar transport (IFT) system for removal from cilia (Desai et al., 2020; Lv et al., 2021).

The post-translational modification of proteins by ubiquitination plays pivotal roles in a wide range of signaling processes (Otten et al., 2021). The covalent attachment of the ubiquitin peptide to a target can change the target’s stability, localization, trafficking, activity, and protein-protein interactions. Ubiquitination requires the activation of ubiquitin by E1 ubiquitin-activating enzymes, the transfer of the ubiquitin to an E2 ubiquitin-conjugating enzyme followed by the ligation of the ubiquitin onto the target protein by an E3 ubiquitin ligase or a complex of E2 and E3 enzymes. There are 2 E1 activating enzymes, ~40 E2 conjugating enzymes and more than 600 E3 ligases encoded in the human genome.

In this work, we examine the role of Arih2 in regulating Smo. Our prior work showed that loss of Arih2 elevated ciliary Smo levels at the basal state and increased the level of Smo in cells (Lv et al., 2021). Arih2, also known as TRIAD1 is a RING-in-between-RING (RBR) type E3 ubiquitin ligase. This ligase has been mostly studied in the context of cancer where it has anti-proliferative effects on myeloid progenitor cells and its expression is reduced in acute myeloid leukemia (Marteijn et al., 2005; Wang et al., 2011). The gene is widely distributed from across metazoans. It is found in organisms like mouse and humans that utilize ciliary Hedgehog signaling, *Drosophila melanogaster* that has non-ciliary Hedgehog signaling, and organisms like *Caenorhabditis elegans* and *Arabidopsis thaliana* that do not have Hedgehog signaling. This distribution suggests that Arih2 regulates other pathways besides Hedgehog (Aguilera et al., 2000; Marín and Ferrús, 2002; Mladek et al., 2003). Consistent with this idea, mice lacking *Arih2* die perinatally with few survivors past the first week of life. Survivors are severely runted and show signs of excessive inflammation (Lin et al., 2013). The authors of this work did not report phenotypes associated with defective Hedgehog signaling however, unpublished work from the International Mouse Phenotyping Consortium documents structural birth defects of the heart, kidney, and skin in heterozygotes that might be Hedgehog related (https://www.mousephenotype.org/data/genes/MGI:1344361). Vertebrate Arih2 undergoes extensive alternative splicing producing nine protein variants in human and two in mouse. We find that in mouse, the isoforms are differentially localized with Arih2α localized to the nucleus and cytoplasm and Arih2β localized to the ER. Only Arih2β rescues the Smo phenotypes in *Arih2*^-/-^ cells suggesting the Arih2β controls the biosynthesis of Smo possibly through ER-associated degradation (ERAD).

## Materials and Methods

### Plasmids

Plasmids were assembled by TEDA method (Xia et al., 2019) into the pHAGE lentiviral backbone (Wilson et al., 2008). All inserts are derived from mouse unless otherwise stated. Mutations were generated by PCR amplification with mutated primers and the corresponding amplicons were TEDA assembled. All inserts were fully sequenced and matched Ensembl reference sequence, NCBI reference sequence, or expected mutant forms. Plasmids are listed in Table S1 and SnapGene files will be provided upon request.

### Cell Culture

Wild-type mouse embryonic fibroblasts (MEFs) were derived from E14 embryos and immortalized with SV40 Large T antigen. These cells were cultured in 95% DMEM (4.5 g/L glucose), 5% fetal bovine serum (FBS), 100 U/mL penicillin, and 100 μg/mL streptomycin (all from Gibco-Invitrogen).

For Smoothened Agonist (SAG) experiments, MEFs were plated at near confluent densities and serum starved (same culture medium described above but with 0.25% FBS) for 24 hr prior to treatment to allow ciliation. SAG (Calbiochem) was used at 400 nM.

Sonic Hedgehog (SHH) conditioned medium was generated from HEK 293T cells expressing HsSHH (BL243), XtScube2 (BL244), and MmDisp1 (BL323). Cells stably secreting SHH were grown to confluency in 90% DMEM (4.5 g/L glucose), 10% FBS, 100 U/mL penicillin, and 100 μg/mL streptomycin, medium was then replaced with low serum medium (0.25% FBS) and grown for 24 hr. Medium was collected, filtered sterilized with 0.45 μm filter (Millipore) and titered for the ability to relocated Smo to cilia. Dilutions similar in effect to 400 nM SAG were used for experiments.

The chemicals used in this study include lysosomal degradation and autophagy inhibitor Bafilomycin A1 (100 nM), protein transport from the endoplasmic reticulum to the Golgi complex inhibitor Brefeldin A (BFA) (50 μg/mL), protein synthesis inhibitor cycloheximide (CHX) (150 μg/mL), the ERAD inhibitor Eeyarestatin I (EerI) (50 μM), proteasomal degradation inhibitor MG132 (1 μM).

### Lentivirus Production

Lentiviral packaged pHAGE-derived plasmids (Wilson et al., 2008) were used for transfection. These vectors are packaged by a third-generation system comprising four distinct packaging vectors (Tat, Rev, Gag/Pol, VSV-g or MLV-env) using HEK 293T cells as the host. DNA (plasmid of interest: 5 μg; Tat: 0.5 μg; Rev: 0.5 μg; Gag/Pol: 0.5 μg; VSV-g/MLV-env: 1 μg) was delivered to the HEK 393T cells as calcium phosphate precipitates. After 48 hr, the supernatant was harvested, filtered through a 0.45 μm filter (Millipore), and added to subconfluent cells. After 24 hr, cells were selected with corresponding antibiotics: nourseothricin (Nat, 50 μg/mL), puromycin (Puro, 1 μg/mL), zeocin (Zeo, 500 μg/mL) or blasticidin (Bsd, 60 μg/mL).

### Flow Cytometry

For flow sorting, pelleted cells were resuspended in the corresponding media and sorted into tubes or 96-well plates containing media with 10% fetal bovine serum by a BD FACS C-Aria II Cell Sorter (BSL-2+/BSC).

### Genome Editing

Guide RNAs were selected from Brie library (Doench et al., 2016) or designed using CHOPCHOP (Labun et al., 2019). Corresponding oligonucleotides were cloned into lentiCRISPR v2 Puro (gift from Feng Zhang [Addgene plasmid # 52961]) (Sanjana et al., 2014) or lentiCRISPR v2 Puro^P93S^ (BL245) and screened by sequencing. lentiCRISPR v2 Puro^P93S^ is similar to its parent except for a proline to serine mutation in the puromycin N-acetyl-transferase gene, which increases its resistance to Puro (https://www.addgene.org). The vectors were packaged into lentiviral particles and transfected into MEFs cells. After selection, the pools were analyzed by flow cytometry. Individual cells were sorted into 96-well plates by flow cytometry or dilution cloning. Single mutant clones were identified with Sanger sequencing, GENEWIZ Amplicon-EZ sequencing, immunofluorescence, or immunoblotting. Sequencing results were analyzed with GEAR Indigo (https://www.gear-genomics.com), Poly peak parser, and SWS method (Hill et al., 2014; Jie et al., 2017; Rausch et al., 2020).

Flag-Avi knock-in was achieved by using CRISPR-Cas9 genome editing. Cas9 and sgRNA were expressed from lentiCRISPR v2 Puro (BL1139). The template for homology-directed repair was designed in Benchling (https://www.benchling.com/). Each homology arm was about 600 bp. For the knock-in experiments, cells were transfected by standard calcium phosphate method or Qiagen Effectene Transfection Reagent according to the manufacturer’s protocol.

### Next Generation Sequencing and Data Analyzing

PCR products were analyzed by Amplicon-EZ paired-end sequencing (Azenta Genewiz). Using bwa-mem2 (https://github.com/bwa-mem2) with default parameters, paired-end sequencing reads from each replicate were first aligned to *Arih2* pseudogenes (*Gm12263* and *Gm49867*), which contain many substitutions compared to the parental *Arih2* gene, unmapped reads were then extracted and mapped onto the enriched region of *Arih2*. Because the cassette exon is specifically spliced out in Arih2β but not Arih2α, sequencing read pairs were deemed to belong to Arih2β if they cover the cassette exon with an overhang of at least 5 nucleotides, otherwise they were considered as reads from Arih2α.

### Immunofluorescence and Live-cell Imaging

Cells were fixed with 2% paraformaldehyde for 15 minutes., permeabilized with 0.1% Triton-X-100 for 2 min and stained as described (Follit et al., 2006). In some cases, fixed cells were treated with 0.05% SDS for 5 min before prehybridization to retrieve antigens. The primary and secondary antibodies are described in Table S2. For thioflavin T fluorescence assay, cells were incubated with 5 μM thioflavin T (Millipore Sigma) for 10 minutes before fixed (Beriault and Werstuck, 2013).

For live-cell imaging, cells were seeded in 35 mm glass bottom, collagen-coated dishes (MatTek Corporation) and cultured for at least 24 hours before imaging.

Confocal images were taken with a LSM910 equipped with a 63X objective and converted to a maximum projection with ZEN 3.1 blue edition (Zeiss).

### Protein and mRNA Analysis

For western blots, cells were pelleted and lysed directly with denaturing gel loading buffer (Tris-HCl 125 mM pH 6.8, glycerol 20% v/v, SDS 4% v/v, β-mercaptoethanol 10% v/v, bromophenol blue). The primary and secondary antibodies are described in table S6. Western blots were developed by chemiluminescence (Super Signal West Dura, Pierce Thermo) and imaged using an Amersham Imager 600 imager (GE Healthcare Life Sciences). Bands were quantified with Gel-Pro Analyzer 4 (Meyer Instruments).

For immunoprecipitations, cells were serum starved for 48 hr and proteins were extracted with lysis buffer (20 mM HEPES pH 7.5, 50 mM KCl, 1mM MgCl_2_) with 0.5% digitonin and protease inhibitor (cOmplete EDTA-Free, Roche). Insoluble components were removed by centrifugation at 20000 g. Primary antibodies pre-adsorbed to protein G-sepharose beads (GE Healthcare) were added to the cell extract and the mixture incubated for 2 hr at 4°C. After centrifugation, beads were washed with lysis buffer supplemented with 0.1% digitonin before elution in denaturing gel loading buffer for SDS-PAGE electrophoresis and Western blotting analysis.

Isolation of mRNA and quantitative mRNA analysis was performed as previously described (Jonassen et al., 2008) using the primers tabulated in Table S3.

### Biotinylation of Cell Surface Proteins

Cell surface proteins were biotinylated by a non-cell permeable EZ-Link Sulfo-NHS-SS-Biotin (Thermo Fisher Scientific) as described (Pusapati et al., 2018). Cells with surface proteins biotinylated were lysed by CelLytic M solution containing cOmplete, EDTA-free Protease Inhibitor Cocktail (Millipore Sigma). The lysate was clarified by centrifugation at 4°C, 18000 g for 10 minutes. Biotinylated proteins were captured and purified from the supernatant by Pierce High Capacity NeutrAvidin Agarose (Thermo Fisher Scientific). After washing six times with lysis buffer, the beads were extracted with denaturing gel loading buffer containing 100 mM DTT at 37°C for 1 hr to release biotinylated proteins.

### Ubiquitination Assay

Smo ubiquitination assay was done as described previously (Lv et al., 2021). In brief, HEK 293T cells were plated at 60% confluent density onto a 10 cm plate. After 24 hours, the cells were transfected with 5 μg of each vector using calcium phosphate transfection. 24 hours after transfection, cells were treated with 50 μM Eeyarestatin I and 1 μM MG132 for 4 hours to block ERAD and proteasomal degradation. Cells were lysed and Smo was captured with anti-Flag M2 affinity gel (Millipore Sigma) as described in the immunoprecipitation method in Protein and mRNA analysis.

### Statistical analysis

Statistical results were obtained from at least three independent experiments. Statistical differences between groups were tested by One-Way ANOVA, Two-Way ANOVA, or repeated measures ANOVA in GraphPad Prism 7.04. Differences between groups were considered statistically significant if *p* < 0.05. Otherwise, non-significant (n.s.) was labeled. Statistical significance is denoted with asterisks (* p < 0.05; ** p < 0.01; *** p < 0.001, **** p < 0.0001). Error bars indicate standard deviation (S.D.).

## Results

### Arih2 regulates cellular and ciliary Smo levels

In a CRISPR-based screen to identify Ub-related genes regulating Hedgehog signaling, we identified the E3 ligase Arih2 as a negative regulator of the pathway whose loss affected Smo levels and ciliary localization (Lv et al., 2021). In our current work, we seek to understand how Arih2 regulates the Hedgehog pathway. Measuring endogenous *Gli1* expression (Figure 1A) reproduced our previous finding with a Hedgehog reporter GreenBomb that the loss of Arih2 elevated basal signaling through the pathway but did not alter induced expression (Lv et al., 2021). As previously shown, loss of Arih2 elevated ciliary Smo levels at the basal state (Figure 1B, C) and increased total cellular Smo levels as detected by western blot (Figure 1Da). Also, the loss of Arih2 does not affect ciliogenesis (Figure S1). The excess Smo appears to be largely in intracellular pools as the amount exposed to the surface in *Arih2* mutant cells was similar to the amount surface-exposed in control cells stimulated with SHH (Figure 1D). Smo mRNA levels are similar in GreenBomb and GreenBomb *Arih2*^-/-^ cells suggesting that the increased Smo results from post-transcriptional mechanisms (Figure 1E).

**Figure 1.**
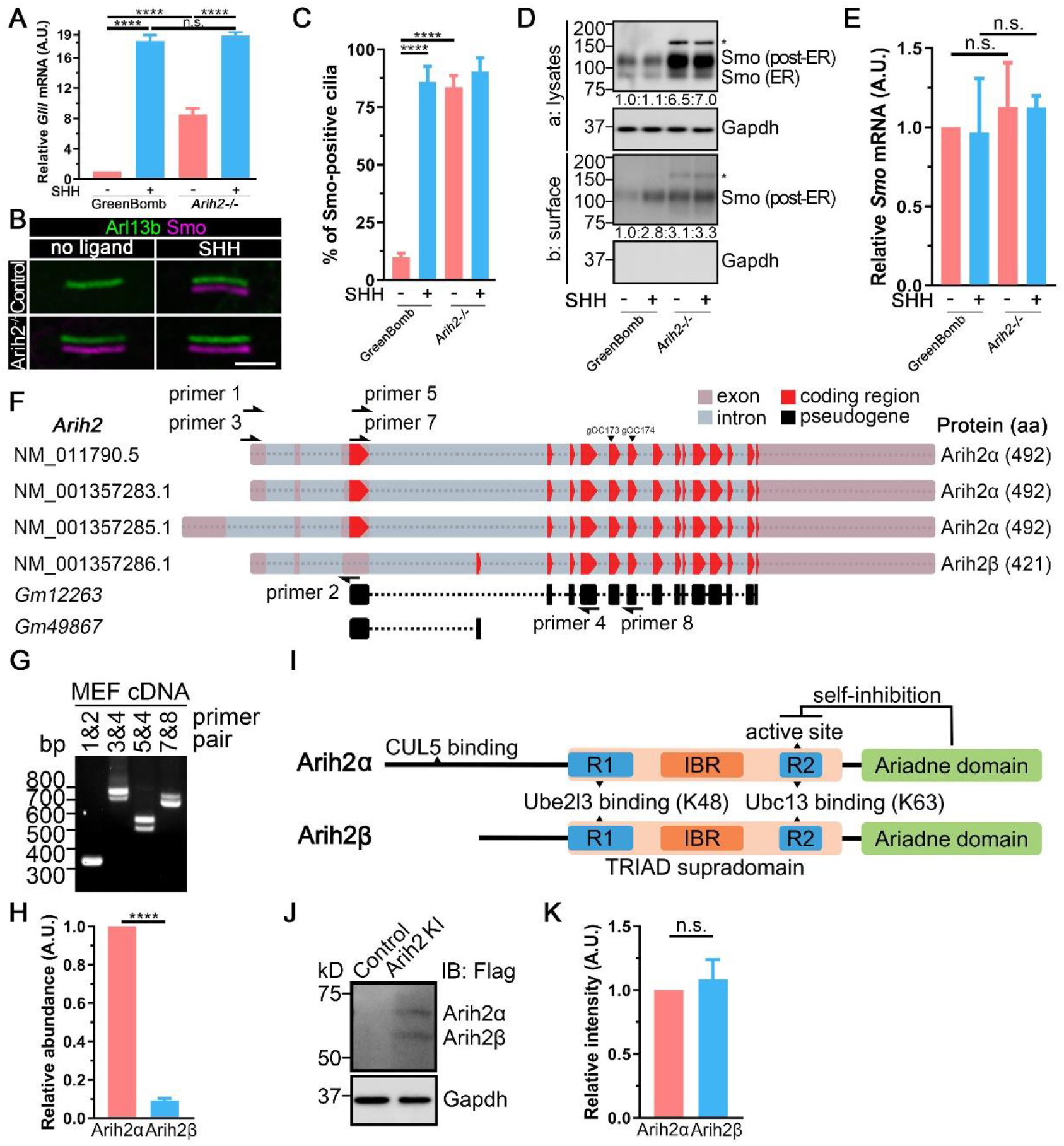
The two isoforms of Arih2 localize differently in cells. **A.** qRT-PCR showing the relative endogenous *Gli1* mRNA in GreenBomb and GreenBomb *Arih2*^-/-^ cells with or without SHH treatment. n = 4 repeats. n.s. not significant, **** p < 0.0001 by Two-Way ANOVA. Error bars indicate SD. **B.** Immunofluorescence showing Smo (Flag, magenta) and cilia (Arl13b, green) in GreenBomb and GreenBomb *Arih2*^-/-^ cells. Scale bar, 3 microns. **C.** Quantitation of Smo-positive cilia described in panel B. n = 6 repeats with 200 cilia counted per experiment. **** p < 0.0001 by Two-Way ANOVA. Error bars indicate SD. **D.** Top (Da) is a western blot of whole-cell extracts from GreenBomb and GreenBomb *Arih2*^-/-^ cells with or without SHH treatment to show total Smo levels. The asterisk marks unspecific bands. Gapdh is a loading control. Bottom (Db) is western blot of surface exposed Smo detected after surface biotinylation and immunoprecipitation. Relative amounts of Smo are listed on the bottom. **E.** qRT-PCR showing the relative *Smo* mRNA in GreenBomb and GreenBomb *Arih2*^-/-^ cells with or without SHH treatment. n = 4 repeats. n.s. not significant by Two-Way ANOVA. Error bars indicate SD. **F.** Diagram of the four *Arih2* splice variants that code for either Arih2α (492 residues) or Arih2β (421 residues). *Gm12263* and *Gm49867* are pseudogenes derived from *Arih2*. **G.** RT-PCR using primers described in F show that MEFs express splice variants that potentially code for both isoforms. **H.** Relative number of reads corresponding to Arih2α and Arih2β by deep sequencing of amplicons with primer set 5&4 against MEF cDNA. **** p < 0.0001 by independent two-sample *t*-test. Error bars indicate SD. **I.** Diagram of the domain structure of Arih2α and Arih2β. R1, RING1; IBR, In-Between-Ring; R2, RING2. R1, IBR and R2 are collectively known as RBR (RING-Between-RING-RING) or TRIAD (two RING fingers and a DRIL [double RING finger linked]). **J.** Western blot of cells with a Flag-tag knocked into *Arih2* just before the stop codon show that approximately equal amounts of each isoform are expressed in MEFs. **K.** Quantification of Arih2 isoform signal intensity of the western blot in I. n = 5 repeats. n.s. not significant by independent samples *t* test. Error bars indicate SD.

### Arih2 has two isoforms

NCBI Gene describes four mouse *Arih2* transcript variants that differ at the N-terminal coding region (Figure 1F). There are also two *Arih2* pseudogenes in the mouse genome, but these are highly mutated and do not appear to express proteins (Figure 1F). The four transcripts encode isoforms of 492 and 421 residues, which we named Arih2α and Arih2β (Figure 1F). Message representing both isoforms is readily detected in fibroblasts (Figure 1G) although Arih2α mRNA levels appear to be about 10-fold higher than Arih2β (Figure 1H). Arih2α and Arih2β are similar with shared ring, ring between ring, and Ariadne domains but differ at their N-termini. The N-terminus of Arih2α carries a cullin-5 binding site that is missing from Arih2β while the N-terminus of Arih2β is predicted to encode a signal peptide with a low probability cleavage site at the end (at the A in position 19) (Figure 1I) (Kelsall et al., 2013). Even though Arih2α is expressed at higher levels than Arih2β, knockin a Flag-tag into the endogenous *Arih2* gene just prior to the stop codon suggests that both isoforms are present in the cell at approximately equal levels (Figure 1J, K).

### Arih2β, but not Arih2α, regulates Smo levels

To ensure that the phenotype in the *Arih2* mutant cells was due to the loss of Arih2, we rescued *Arih2* mutant cells with constructs that express Myc-tagged Arih2α or Arih2β, and Myc-tagged Arih2α^C309A^ or Arih2β^C238A^ in which the active sites are mutated (Figure 2A). Interestingly expression of Arih2β returned ciliary Smo (Figure 2B, C) and total Smo (Figure 2D) back to normal while expression of Arih2α was not effective. Arih2β^C238A^ was not functional indicating that the E3 ubiquitin ligase activity of Arih2β is required for rescue (Figure 2B-D).

**Figure 2.**
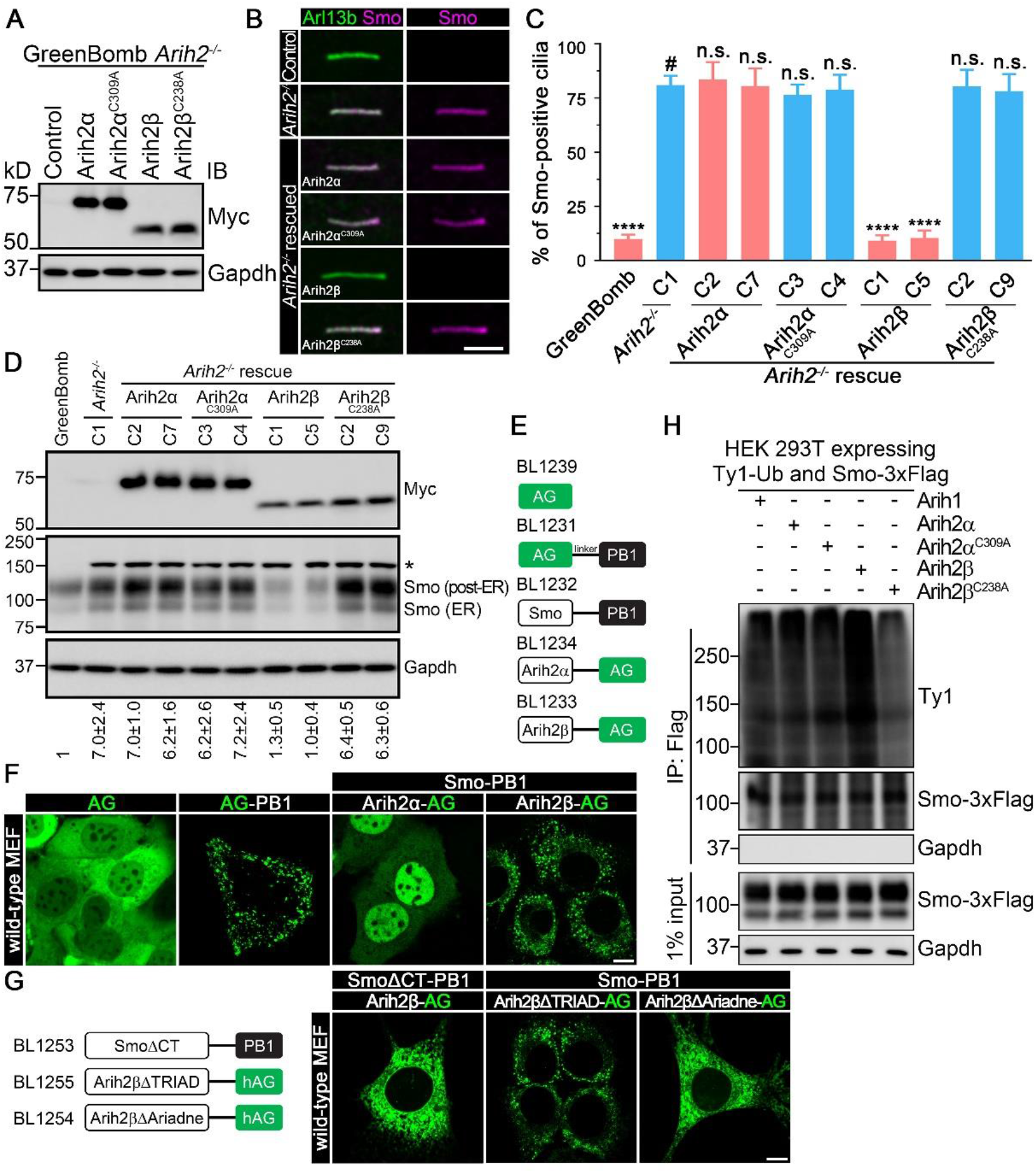
Arih2β regulates Smo level and location. **A.** Western blot of *Arih2*^-/-^ cells (control) and *Arih2*^-/-^ cells rescued with Myc-tagged Arih2α (BL256) and Arih2β (BL929) along with the enzymatic dead versions Arih2α^C309A^ (BL368) and Arih3β^C238A^ (BL930). **B.** Immunofluorescence showing Smo (Flag, magenta) and cilia (Arl13b, green) in MEF^Smo-3xFlag^ (control), *Arih2* knockout cells and cells rescued with Myc-tagged Arih2α and Arih2β along with the enzymatic dead versions Arih2α^C309A^ and Arih3β^C238A^. Scale bar, 3 microns. **C.** Quantification of ciliary Smo localization in B. n = 6 repeats with 200 cilia counted per experiment. n.s. not significant, **** p < 0.0001 by Two-Way ANOVA compared to *Arih2*^-/-^ cells (labeled with #). Error bars indicate SD. **D.** Western blots to quantitate rescue of total Smo levels by Myc-tagged Arih2α and Arih2β along with the enzymatic dead versions Arih2α^C309A^ and Arih2β^C238A^. Two independent rescue lines are shown for each construct. The amount of total Smo relative to wild type (GreenBomb) is shown on the bottom. The asterisks mark unspecific bands. Gapdh is a loading control. **E.** Diagram of Fluoppi constructs used. The AG domain is an azami-green fluorescent reporter and the PB1 domain consists of homodimerization domain from the p62 autophagy protein. Fluorescent puncta appear in the cytoplasm when the two domains are brought together as fusion protein (AG-PB1 positive control) or by a protein-protein interaction (Watanabe et al., 2017). **F.** Live cell images of MEF cells expressing each of the constructs. Note that cells expressing Smo-PB1 with Arih2β-AG show cytoplasmic puncta similar to the positive control AG-PB1. Scale bar, 5 microns. **G.** Live cell images of MEF cells expressing Smo deleted of the C-terminal tail (BL1253) and Arih2β deleted of the TRIAD domain (BL1255) or Ariadne domain (BL1254). Scale bar, 5 microns. **H.** Ligation of Ub onto Smo by Arih2. HEK 293T cells expressing Smo-3xFlag (PD22) and Ty1-Ub (BL1035) were transiently transfected with the constructs indicated on the top. After lysis, Smo was immunoprecipitated with Flag resin and examined by western blot with antibodies listed on the right side. Input are extracts before Flag immunoprecipitation. A lighter exposure of the Ty1 western blot is included.

To determine if Arih2 is capable of ubiquitinating Smo, we transfected HEK293 cells with Smo-Flag and Ty1-Ub along with either Arih1, Arih2α, Arih2α^C309A^, Arih2β, or Arih2β^C238A^. Immunoprecipitating Smo-Flag and measuring the incorporation of Ub-Ty1 indicates that cells expressing Arih2β incorporated more Ub onto Smo than cells expressing the Arih2β active site mutation or the Arih2α and Arih1 isoforms (Figure 2H).

To explore physical interactions between Smo and Arih2, we used the recently developed fluorescent protein-protein interaction (Fluoppi) approach (Watanabe et al., 2017). In this method, one protein is tagged with an azami-green (AG) fluorescent tag and the other protein is tagged with the PB1 homodimerization domain from the p62 autophagy protein (Figure 2E). If the bait and prey proteins do not interact the fluorescence will be dispersed throughout the cell but if the proteins interact, fluorescent puncta appear in the cytoplasm. Co expressing Arih2α-AG with Smo-PB1 showed dispersed fluorescence indicating no interaction. However, a co expression of Arih2β-AG with Smo-PB1 showed strong puncta indicating a physical interaction (Figure 2F). Deletion of the Ariadne domain of Arih2β or the C-terminal tail of Smo abolished the interaction while the interaction was maintained when the Triad domain was deleted (Figure 2G).

### Arih2β localizes in the ER

Previously we reported that Arih2 was localized predominately in the nucleus with a lesser amount in the cytoplasm and none was detected in the cilium (Lv et al., 2021). However, this work only examined the 492-residue Arih2α isoform. Since the Arih2β isoform is the relevant isoform, we repeated this work and examined both isoforms. We could not detect ciliary localization with either isoform (Figure 3A). As previously shown, Arih2α localizes predominately in the nucleus with some cytoplasmic localization in MEFs (Figure 3A, B), which is similar to the distribution of its paralog Arih1 (Figure S2A). Arih2α in the cytoplasm did not appear to associate with any cytoskeletal or vesicular structures. In contrast to the prominent nuclear localization of Arih2α, Arih2β localizes in cytoplasm with no localization to the nucleus (Figure 3B). Within the cytoplasm, Arih2β was concentrated near the nucleus where it associated with tubular and vesicular structures, and colocalized with cell body Smo (Figure 3B). Similar localization of both isoforms was seen in IMCD3, hTERT RPE-1, HEK 293T and NIH/3T3 cells (Figure S2B). No association was seen between Arih2β and endosomes or the Golgi complex (Figure S2C, D), and the localization of Arih2β was not affected by Brefeldin A (BFA) treatment (Figure S2D). However, extensive co-localization was observed between Arih2β and an ER-targeted GFP construct or the ER localized ubiquitin E3 ligase Syvn1 (Figure 3C) (Kaneko et al., 2002).

**Figure 3.**
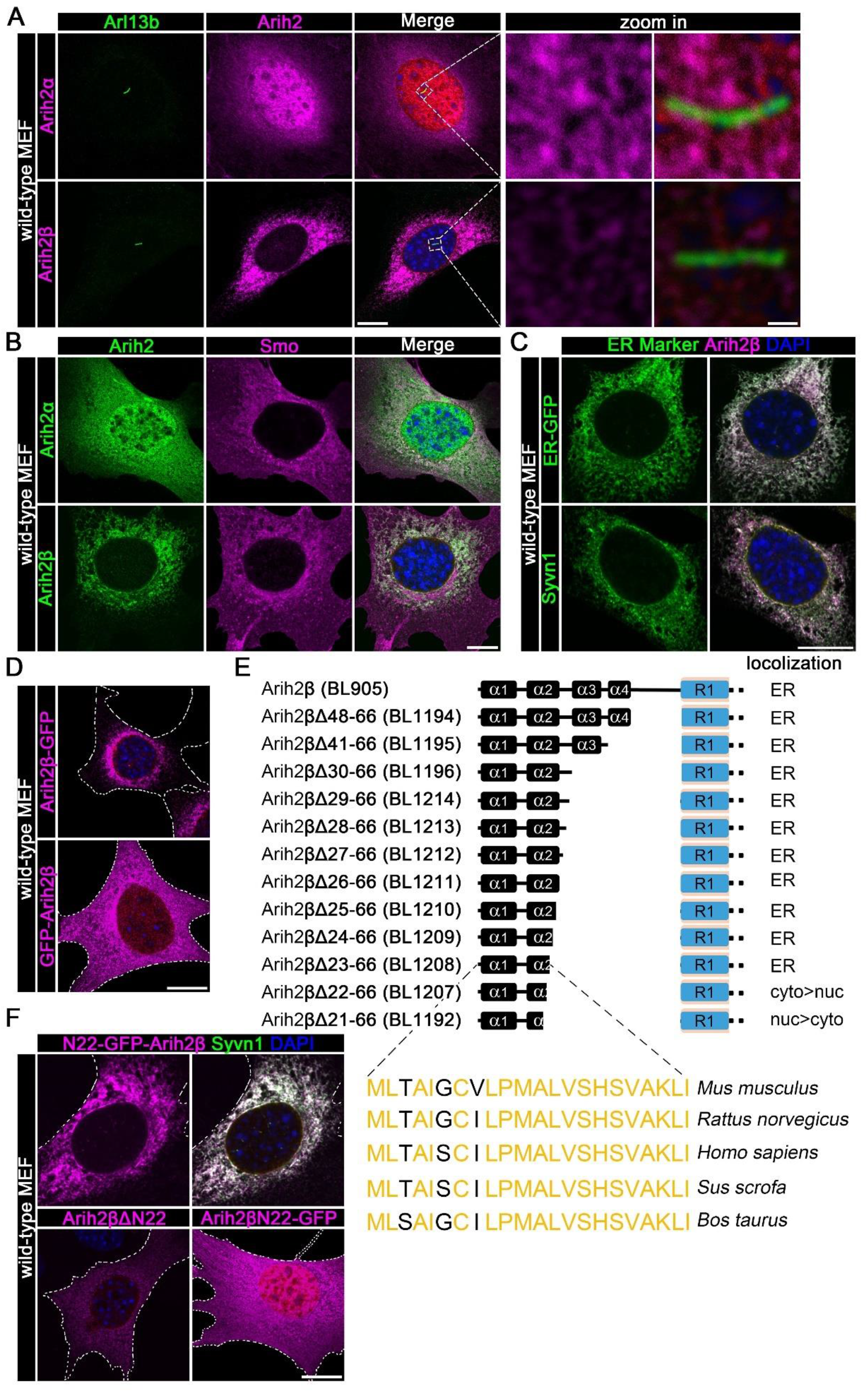
Arih2β localizes to the endoplasmic reticulum. **A.** Wild-type MEF cells expressing Flag-tagged Arih2α (BL225) and Arih2β (BL905) were stained for Arih2 (Flag, magenta), cilia (Arl13b, green), and DNA (DAPI, blue). Scale bar, 5 microns in original images and 1 micron in enlarged images. **B.** Wild-type MEF cells expressing Flag-tagged Arih2α (BL225) and Arih2β (BL905) were stained for Arih2 (Flag, green), Smo (magenta), and DNA (DAPI, blue). Scale bar, 5 microns. **C.** Wild-type MEF cells expressing Flag-tagged Arih2β and the ER markers ER-GFP (BL1099) or Syvn1-HA (BL1087) were stained for Arih2 (Flag, magenta), the ER marker (GFP or HA, green), and DNA (DAPI, blue). Scale bar, 5 microns. **D.** Wild-type MEF cells expressing Arih2β-GFP (BL1117) and GFP-Arih2β (BL1118). Cell outlines are shown by dotted lines. Scale bar, 5 microns. **E.** Diagram of the N terminus of Arih2β with a series of truncations produced to test the possible ER signal peptide. The four alpha helixes were predicted by PSIPRED 4.0. **F.** Wild-type MEF cells expressing Arih2β with GFP inserted after residue 22 (N22-GFP-Arih2β, BL1218, magenta) and Syvn1-HA (BL1087, green). Bottom row: Wild-type MEF cells expressing Arih2β missing the first 22 residues (Arih2βΔN22, BL1217, magenta) and the first 22 residues of Arih2β fused to GFP (Arih2βN22-GFP, BL1219, magenta). Scale bar, 5 microns.

Arih2α and Arih2β differ at their N-termini. Arih2β’s N-terminus is critical for localizing it to the ER as C-terminal GFP fusion is retained at the ER while an N-terminal GFP fusion is dispersed throughout the cell (Figure 3D). To identify the ER localization signal of Arih2β, we generated a series of deletions where the N-terminal helical domains were progressively removed starting at the RING1 domain and moving back toward the N-terminus. Immunofluorescence results showed that the first 22 residues, which are evolutionally conserved in mammals, are necessary for the ER localization of Arih2β (Figure 3E). Inserting GFP after the 22^nd^ residue did not disrupt ER localization. However, GFP fusions that carry only the first 22 Arih2β residues are not as highly enriched at the ER suggesting that other portions of Arih2β also contribute to ER localization (Figure 3F). Fluoppi analysis indicated that the Ariadne domain was needed for interaction with Smo (Figure 2G) indicating that Smo binding might enhance ER enrichment of Arih2β.

### Arih2β localizes in the cytoplasmic face of the ER and interacts with Smo

The signal sequence at the N-terminus of Arih2β could function to tether Arih2β on the outer surface of the ER or it could direct the translocation of Arih2β into the lumen of the ER. There is currently no evidence for ubiquitination activity in the lumen of the ER suggesting that the signal sequence is more likely to function to tether the protein on the intracellular surface of the ER. However, to distinguish these possibilities, we made Arih2β fusions to the calcium-measuring organelle-entrapped protein indicators (CEPIA) (Suzuki et al., 2014) (Figure 4A, B). CEPIA is calcium indicator that is fluorescent when exposed to high calcium levels as found in the ER lumen but is non-fluorescent in the lower calcium environment of the cytoplasm (Figure 4C). Targeting CEPIA to the ER by fusion with a modified ER signal sequence from the mouse immunoglobulin heavy chain variable region yielded high fluorescent signal. However, fusion of CEPIA to C terminus of Arih2β or inserting it after the signal sequence at the N-terminus correctly localized the indicator to the ER but produced little signal (Figure 4C). The lack of signal indicates that the CEPIA domain in these constructs is in the cytoplasm supporting a model where the signal sequence of Arih2β tethers the protein onto the cytoplasmic face of the ER.

**Figure 4.**
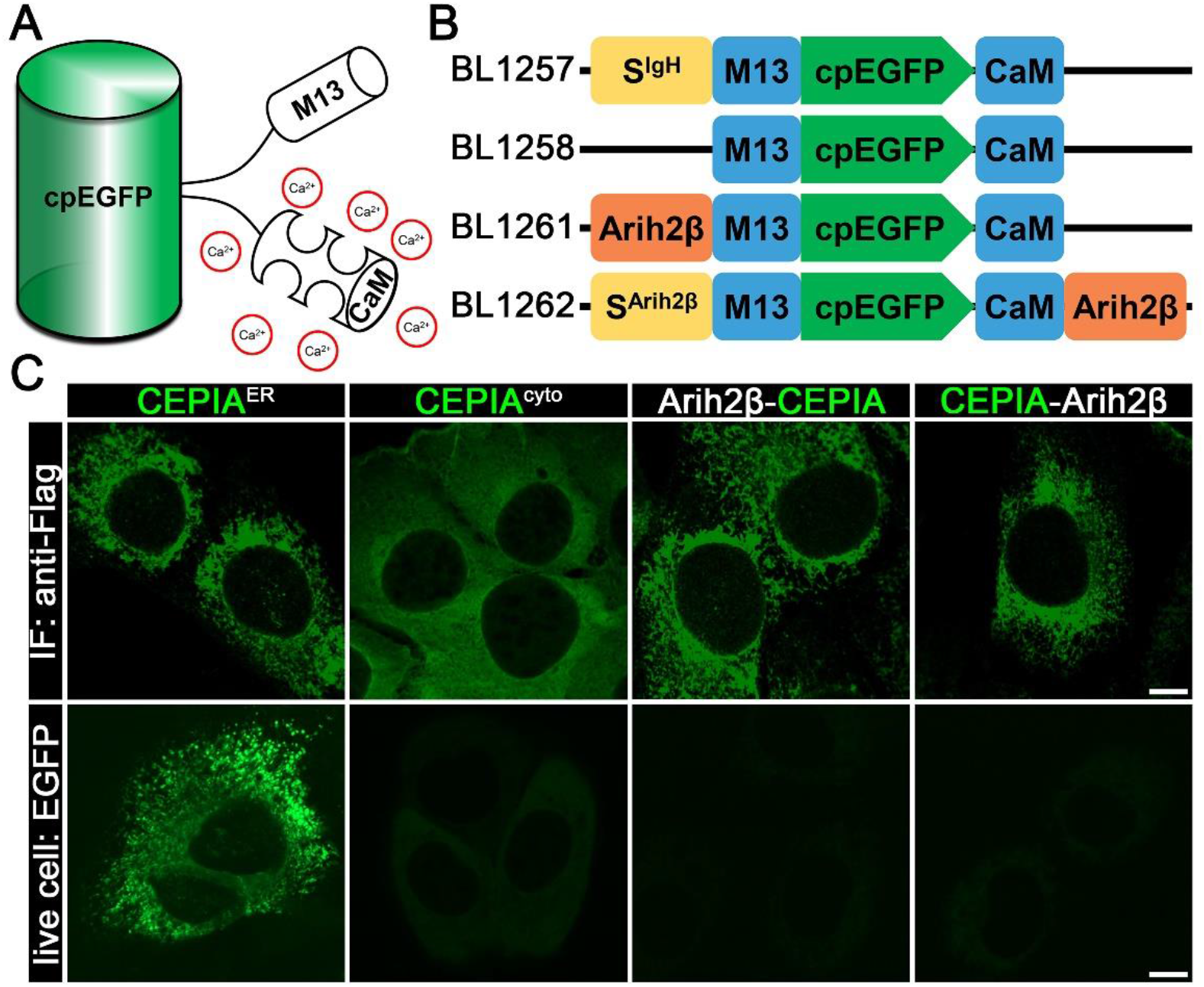
Arih2β localizes in the cytoplasmic side of the ER. **A.** Diagram of the CEPIA calcium indicator, which is a circularly permutated EGFP (cpEGFP) flanked by the myosin light chain M13 helix and calmodulin (CaM). **B.** Diagram of constructs used. S^Arih2β^ is the first 22 residues of Arih2β. **C.** MEFS expressing each of the constructs in B was stained for Flag (top row) or imaged live for CEPIA fluorescence (bottom row). Scale bars, 5 microns.

### The ubiquitin-proteasome system (UPS) is the major pathway for the degradation of Smo

Our finding that the loss of the ER localized isoform of Arih2 causes Smo protein levels to increase without increasing Smo transcription suggests that Arih2β is involved in regulating either the biosynthesis or degradation of Smo. To examine the turnover of Smo, we analyzed protein stability by measuring Smo levels after stopping biosynthesis with cycloheximide (CHX). Smo was degraded over a 12-hour period alone with a half-life of 4 hours (Figure 5A and F). Degradation of Smo was inhibited by both the lysosome inhibitor BafA1 and the proteasome inhibitor MG132 increasing the half-lives to 6 hours and 9.8 hours, respectively (Figure 5B, C, F). These findings suggest that Smo can be degraded by both the proteasome and lysosome.

**Figure 5.**
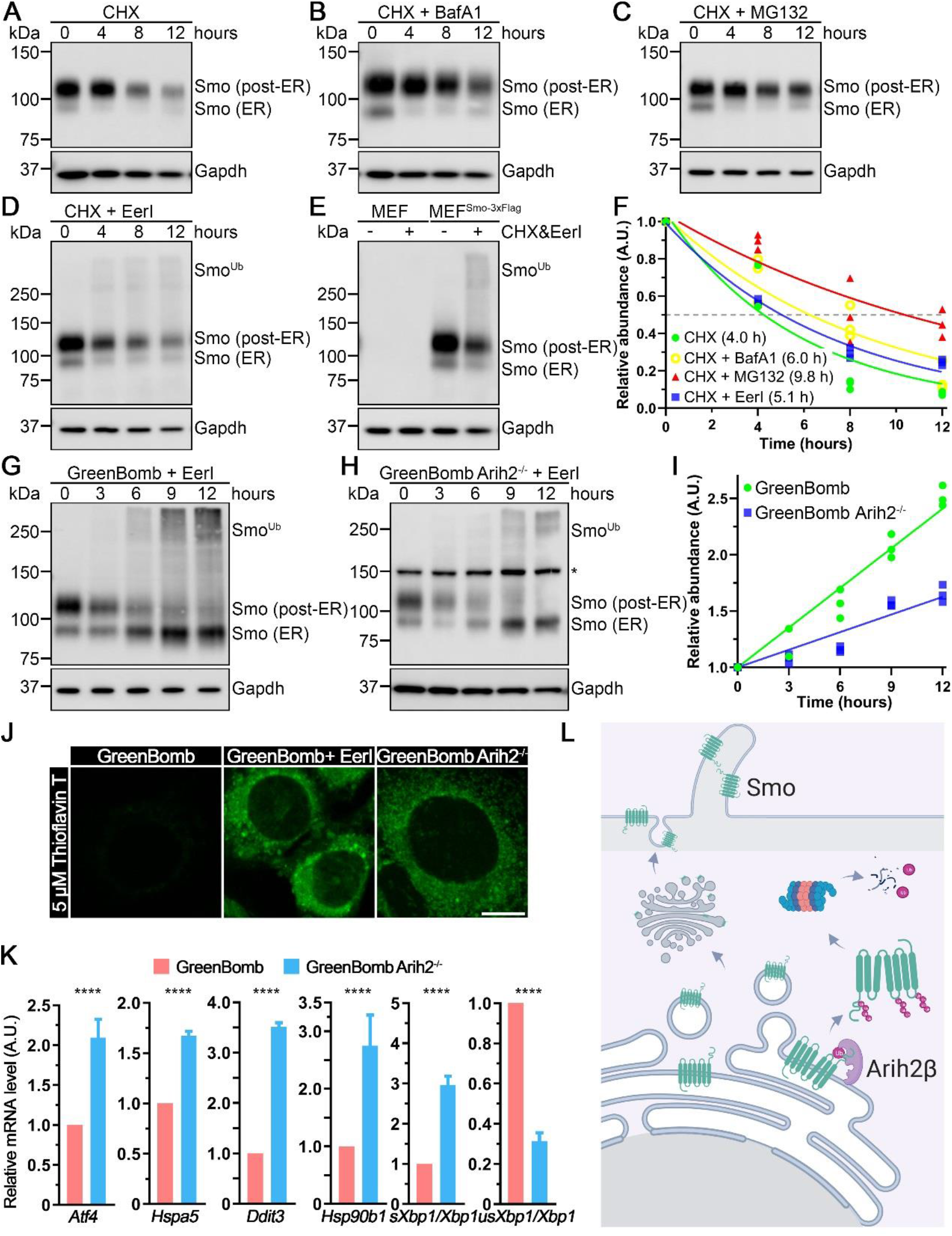
Arih2β mediates Smo ubiquitination and ERAD. **A-D.** MEF^Smo-3xFlag^ were treated with cycloheximide (CHX) to stop new protein synthesis and no drug (**A**), BafA1 to inhibit autophagy and the lysosome (**B**), MG132 to inhibit the proteosome (**C**), or EerI to inhibit ERAD (**D**). Cells were collected at 0, 4, 8 and 12 hrs and analyzed by anti-Flag western blot. Gapdh is a loading control. **E.** Anti-Flag western blot of MEFs and MEF^Smo-3xFlag^ either untreated or treated with CHX and EerI. Note large ubiquitinated forms of Smo when treated with CHX and EerI. **F.** Half-lives of Smo under the conditions in **A-D**. ER, post-ER, and ubiquitinated Smo were combined for this analysis. N=3. The exponential decay equation model used is Y = (Y0-Plateau)*exp(-K*X)+Plateau. The goodness of fit described by R squared for A-D is 0.8962, 0.9154, 0.8210, and 0.9858, respectively. **G-H.** GreenBomb (**G**) and GreenBomb *Arih2*^-/-^ (**H**) cells were treated with EerI but not CHX. Cells were collected at 0, 4, 8 and 12 hrs and analyzed by anti-Flag western blot. **I.** Quantification of the ubiquitinated forms of Smo (signal above * at 150 kD) in **G-H**. N=3. Linear curve fitting is used here. The slopes of linear regression equations established differ significantly (p < 0.001). **J.** Thioflavin T fluorescence staining of aggregated proteins in GreenBomb, GreenBomb treated with EerI, and GreenBomb *Arih2*^-/-^. Scale bar is 5 microns. **K.** qRT-PCR showing the relative endogenous *Atf4*, *Hspa5*, *Ddit3*, *Hsp90b1*, *sXbp1/Xbp1*, and *usXbp1/Xbp1* mRNA in GreenBomb and GreenBomb *Arih2*^-/-^ cells. n = 3 repeats. **** p < 0.0001 by independent samples *t* test. Error bars indicate SD. **L.** Model for the function of Arih2 in regulating Smo levels in the cell. ER-localized Arih2β recognizes misfolded Smo and ubiquitinates cytoplasmic lysine residues. The ubiquitinated Smo is extracted from the membrane and sent to the proteosome for degradation. Normally folded Smo exits the ER and traverses the Golgi complex before delivery to the plasma membrane and cilium.

In general, transmembrane proteins are degraded through the lysosomal pathway, but misfolded transmembrane proteins can undergo proteasomal degradation via ER-associated degradation (ERAD). During ERAD, cytoplasmic domains of misfolded membrane proteins become ubiquitinated targeting them for extraction from the membrane by p97/Cdc48 and degradation by the proteosome (Bodnar and Rapoport, 2017). Treatment with Eeyarestatin I (EerI), which blocks ERAD by inhibiting p97/Cdc48 (Wang et al., 2008), attenuated the degradation of Smo (Figure 5D) and extended the half-life to 5.1 hrs. Unlike treatment with CHX or MG132, treatment with EerI produced a smear of higher molecular weight forms of Smo. These higher molecular weight forms are not seen in the parental cell line indicating that they are not a cross reactive product (Figure 5E). It is likely that these larger forms are poly ubiquitinated forms of Smo that under normal conditions would be degraded by ERAD but accumulate when ERAD is blocked. If Arih2 responsible for targeting misfolded Smo for degradation, we expect that the amount of ubiquitinated Smo that accumulates with EerI treatment should be reduced in Arih2 mutants. Supporting this idea, blocking ERAD with EerI without blocking translation with CHX caused a greater accumulation of ubiquitinated Smo in control cells compared to Arih2 mutant cells (Figure 5G, H, I).

Our finding that ER-localized Arih2β regulates the total cellular levels of Smo suggests that Arih2β functions as a quality control sensor in the ER targeting misfolded Smo for ERAD. The ER accumulation of Smo and other potential Arih2β substrates in *Arih2* mutants is likely to cause ER stress and the unfolded protein response. Consistent with this idea, protein aggregates as detected with thioflavin T (Beriault and Werstuck, 2013) were as abundant in *Arih2* mutant cells as they are in control cells treated with the ERAD inhibitor EerI (Figure 5J). As a more direct test, we measured expression of unfolded protein response target genes (Oslowski and Urano, 2011). Changes were consistent with increased UPR with *Atf4*, *Hspa5*, *Ddit3*, *Hsp90b1* and spliced *Xbp1* increased and unspliced *Xbp1* decreased in *Arih2* mutant cells (Figure 5K).

## Discussion

Our findings show that Arih2 loss elevates basal expression of Hedgehog responsive genes, increases the total cellular level of Smo and causes Smo to accumulate in cilia at the basal state. Arih2, also known as Triad1 is a ring-between-ring E3 ligase that functions with the E2 conjugating enzyme Ube2l3 to ubiquitinate substrates. Consistent with Ube2l3 being a functional E2 for Arih2, we found that Ube2l3 loss also elevated cellular levels of Smo (Lv et al., 2021). In mouse, Arih2 encodes two major isoforms that differ at their N-termini. Arih2α has a longer N-terminus that includes a cullin-5 binding site and localizes to the nucleus while the N-terminus of Arih2β has hydrophobic helix that anchors it to the cytoplasmic face of the ER. The ER-localized Arih2β isoform fully rescues the Smo phenotypes while the nuclear-localized form does not. Most work on Arih2 focuses on Arih2α function as part of the cullin-5 complex (Hüttenhain et al., 2019; Kelsall et al., 2013; Kostrhon et al., 2021). Complete loss of Arih2 in mouse leads to embryonic lethality and increased inflammatory responses (Lin et al., 2013). Heterozygotes show a variety of cilia-related phenotypes including polycystic kidney disease and structural birth defects in kidney, skin, bone, and heart (https://www.mousephenotype.org/data/genes/MGI:1344361) (Bult et al., 2019; Dickinson et al., 2016).

The localization of Arih2β at the ER and our finding that loss of Arih2 elevates total Smo levels suggests that Arih2 regulates the production of Smo, possibly through a quality control mechanism (Figure 5L). Misfolded proteins in the ER are targeted for degradation by the ERAD system. During ERAD, misfolded proteins are targeted to a dislocation complex in the ER membrane where the misfolded protein is polyubiquitinated on cytoplasmic lysine residues. The polyubiquitinated protein is removed from the membrane by the p97/Cdc48 complex and sent to the proteosome for degradation. The major E3 ligases involved in ERAD are thought to be Syvn1 and Amfr. However, more than a dozen other E3s have been implicated in ERAD of specific substrates (Olzmann et al., 2013) and more than 25 E3s localize to the ER (Fenech et al., 2020). Our finding that loss of Arih2 reduces the incorporation of Ub onto Smo when ERAD is blocked by the p97/Cdc48 inhibitor EerI supports a role for Arih2β in ERAD. However, Arih2 is thought to catalyze only the initial mono-ubiquitination (Hüttenhain et al., 2019; Kelsall et al., 2013) indicating that additional E3 ligases are needed to extend the chain. Syvn1 is an interesting possibility as its loss also elevates expression of Hedgehog responsive genes, but we did not observe increased ciliary Smo levels in the knockouts (Lv et al., 2021).

The loss of Arih2 elevates the total level of Smo in the cell. The majority of the extra Smo is not exposed on the cell surface. However, exposed Smo is increased in the mutant to about the level seen in activated control cells. Smo is thought to constantly diffuse into cilia with regulated removal dictating the ciliary level (Milenkovic et al., 2009). Increased plasma membrane Smo would increase the amount that diffuses into the cilium. Over expression of Smo is sufficient to saturate the retrieval process (Corbit et al., 2005) and so it is likely that this is the reason for elevated ciliary Smo in Arih2 mutants. However, ciliary localization of Smo is not sufficient to activate the pathway raising the question of why the loss of Arih2 elevates basal expression of Hedgehog responsive genes. Activation of Smo is driven by phosphorylation and sterol binding (Deshpande et al., 2019; Jia et al., 2004; Nedelcu et al., 2013; Zhang et al., 2004) but point mutations such as the SmoM2 W539L mutation activate the pathway independent of upstream signals. This mutation is thought to shift helixes changing the protein into active conformation (Huang et al., 2018). It is possible that some misfolded protein that escapes the ER in the absence of Arih2 is in an active conformation analogous to the SmoM2 state. Another possibility is that if the loss of Arih2 causes Smo to remain in the ER for an extended time, the high Ca^2+^ environment of the ER lumen may promote the esterification of Smo with cholesterol and drive it towards an active conformation (Hu et al., 2022).

In summary, our work identifies a new regulatory mechanism in the ER that controls the cellular levels of Smo. When this mechanism is defective, ciliary Smo levels are elevated and basal expression through the pathway is increased.

## Abbreviations

IFT: intraflagellar transport
Ub: ubiquitin
Mm: *Mus musculus*
Hs: *Homo sapiens*
Gg: *Gallus gallus*
GPCR: G protein-coupled receptor
DAPI: 4′,6-diamidino-2-phenylindole
SAG: Smoothened Agonist
SHH: sonic hedgehog
NLS: nuclear localization sequence
Bsd: blasticidin
Puro: puromycin
Nat: nourseothricin
Zeo: zeocin
Xt: *Xenopus tropicalis*
NLS: nuclear localization sequence
BFA: Brefeldin A
CHX: cycloheximide
ERAD: ER-associated degradation
Fluoppi: fluorescent protein-protein interaction
AG: azami-green
cpEGFP: circularly permutated EGFP
CaM: calmodulin (CaM)

## Acknowledgments

We thank Dr. Carol E. Schrader and the staff of the University of Massachusetts Medical School Flow Cytometry Core for assistance during this project.

## Author Contributions

Conceptualization: B.L. and G.J.P.; Methodology: B.L., X-O.Z. and G.J.P.; Validation Verification: B.L. and G.J.P.; Formal Analysis: B.L., X-O.Z. and G.J.P.; Investigation: B.L. and G.J.P.; Writing – Original Draft Preparation: B.L. and G.J.P.; Writing – Review & Editing: B.L., X-O.Z. and G.J.P.; Visualization Preparation: B.L.; Supervision: G.J.P.; Funding Acquisition: G.J.P.

## Data and materials availability

All data is available in the main text or the supplementary materials.

## Supplementary Materials

Materials and Methods

Figures S1–S2

Tables S1–S3

References

---

**Figure S1.**
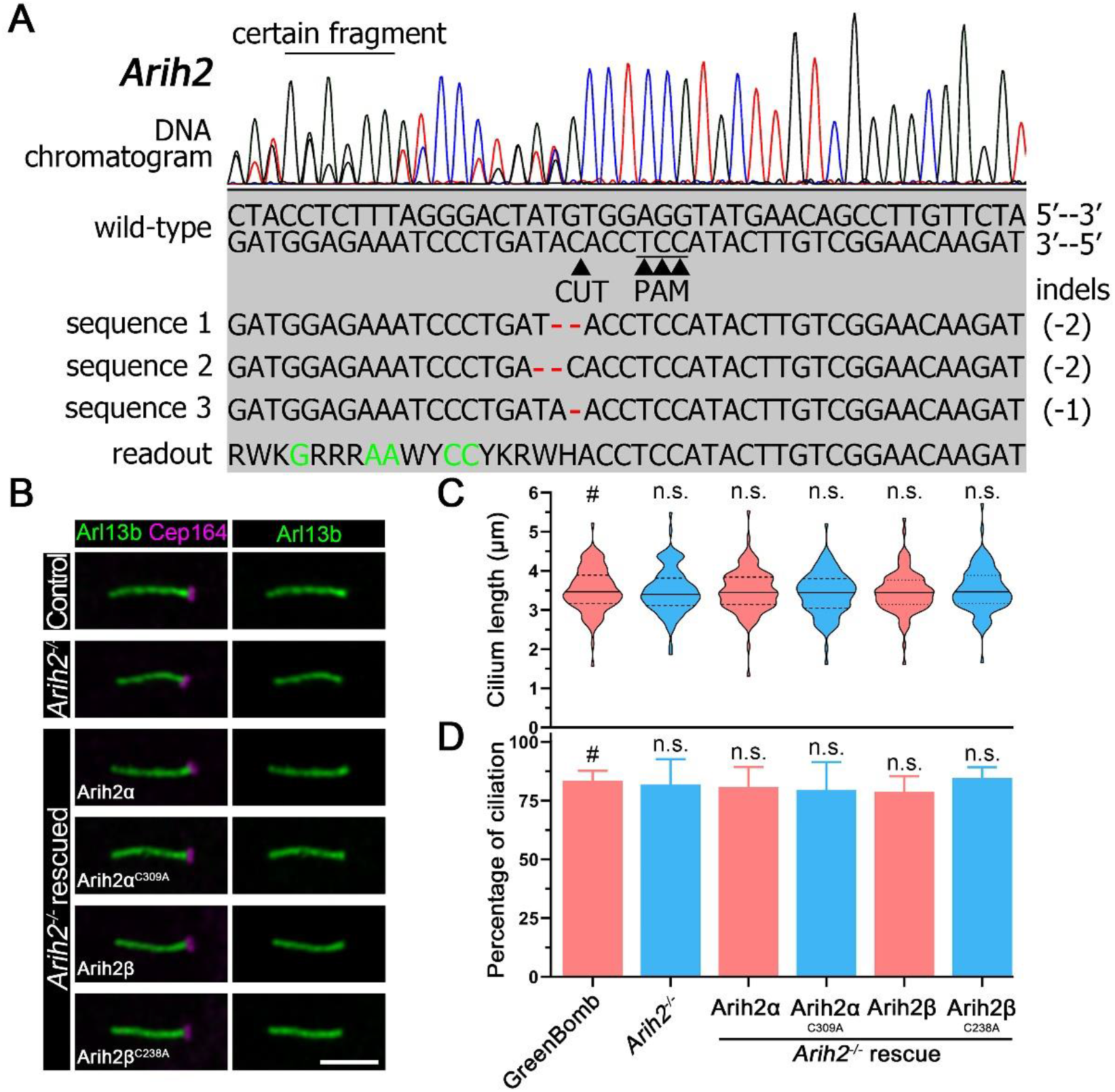
Arih2 knockout doesn’t affect ciliogenesis. **A.** Chromatogram showing one typical clone (gOC173 C1) of GreenBomb *Arih2*^-/-^ cell with deconvolved sequence. Red dashes mark deletions. Green letters in the sequence means defined bases in the readout. **B.** Immunofluorescence of control, *Arih2*^-/-^ and *Arih2*^-/-^ cells rescued with Myc-tagged Arih2α and Arih2β along with the enzymatic dead versions Arih2α^C309A^ and Arih3β^C238A^ stained for cilia (Arl13b, green) and basal bodies (Cep164, magenta). Scale bar, 3 microns. **C.** Quantification of ciliary length in *Arih2*^-/-^ and rescue cells. n = 100 repeats. n.s., not significant by One-Way ANOVA as compared to control (GreenBomb, labeled with #). **D.** Quantification of percent ciliation in *Arih2*^-/-^ and rescue cells. n = 5 repeats. n.s., not significant by One-Way ANOVA as compared to control (GreenBomb, labeled with #).

**Figure S2.**
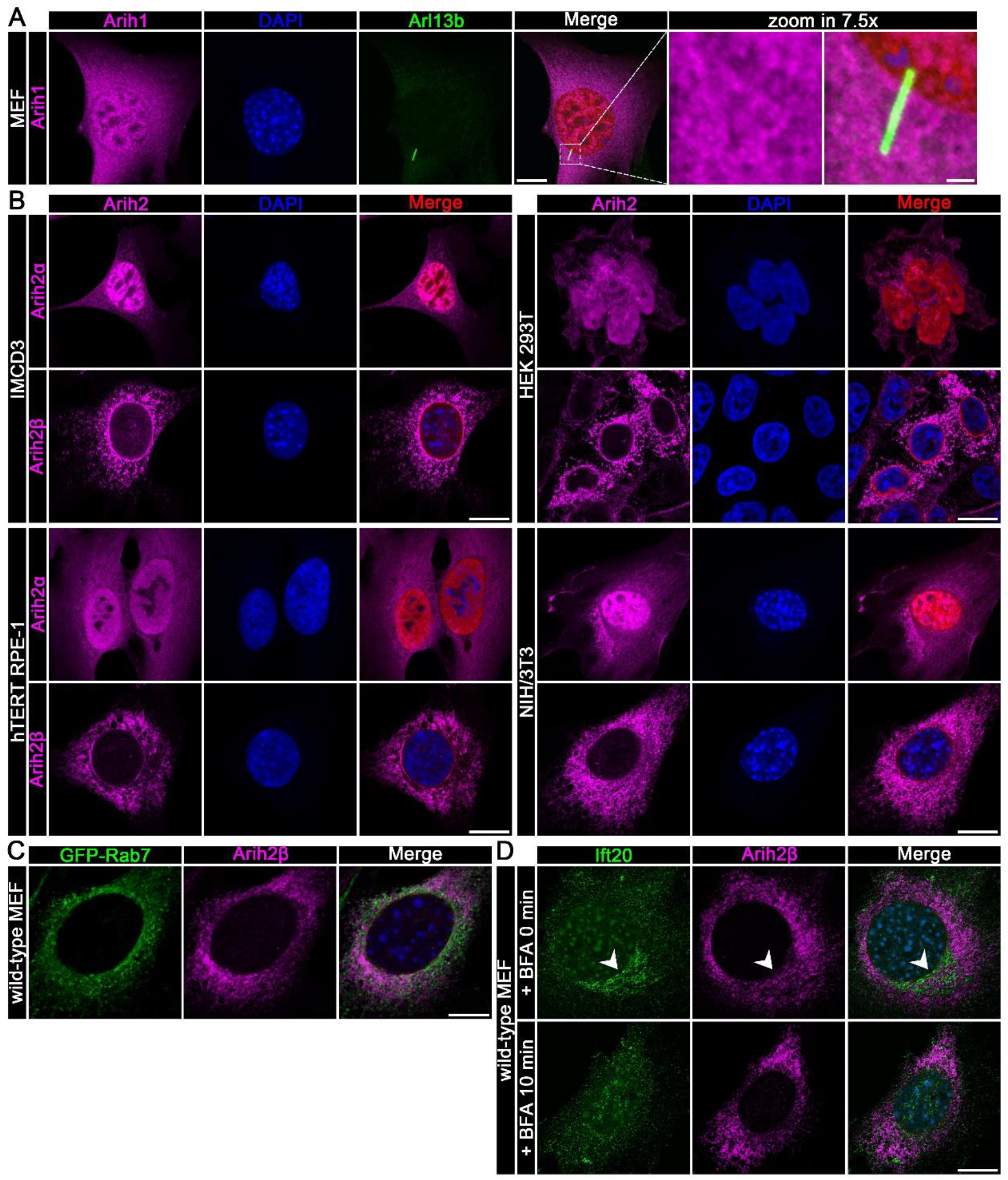
Arih2α and Arih2β localization in diverse cell lines. **A.** Wild-type MEF cells expressing Flag-tagged Arih1 (BL1046) were stained for Arih1 (Flag, magenta), cilia (Arl13b, green), and DNA (DAPI, blue). Scale bar, 5 microns in original images and 1 micron in enlarged images. **B.** IMCD3, HEK 293T, hTERT RPE-1, and NIH/3T3 cells transfected with Flag-tagged Arih2α (BL225) and Arih2β (BL905) and stained for Arih2 (Flag, magenta) and DNA (DAPI, blue). Scale bar, 5 microns. **C.** Wild-type MEF cells expressing Flag-tagged Arih2β (BL905) and GFP-Rab7 (BL1086) stained for Arih2 (Flag, magenta) and GFP (green). Scale bar, 5 microns. **D.** Wild-type MEF cells expressing Flag-tagged Arih2β (BL905) were stained for Arih2 (Flag, magenta) and Ift20 (green) before and after treatment with BFA. Note that BFA disperses the Ift20 pool but not the Arih2β pool. Scale bar, 5 microns.

**Table S1.** Plasmids.

| **Name** | **Description** | **Promoter** | **Tags** | **Drug** |
| --- | --- | --- | --- | --- |
| #52961 | lentiCRISPR v2 Puro | EF1α; U6 | Flag | Puro |
| #58215 | pCMV G-CEPIA1er | CMV | Myc | Neo |
| BL225 | Arih2α-3xFlag (492aa) | EF1α | Flag | Bsd |
| BL243 | HsSHH | CMV | None | Puro |
| BL244 | XtScube2-3xFlag | EF1α | Flag | Bsd |
| BL245 | lentiCRISPR v2 Puro<sup>P93S</sup> | EF1α; U6 | Flag | Puro<sup>P93S</sup> |
| BL256 | Arih2α-6xMyc (492aa) | CMV | Myc | Zeo |
| BL323 | MmDisp1-6xMyc | CMV | Myc | Zeo |
| BL368 | Arih2α<sup>C309A</sup>-6xMyc (492aa) | CMV | Myc | Zeo |
| BL905 | Arih2β-3xFlag (421aa) | EF1α | Flag | Bsd |
| BL929 | Arih2β-6xMyc (421aa) | CMV | Myc | Puro |
| BL930 | Arih2β<sup>C238A</sup>-6xMyc (421aa) | CMV | Myc | Puro |
| BL1035 | 3xTy1-Ub | CMV | Ty1 | Neo |
| BL1046 | Arih1-3xFlag | EF1α | Flag | Bsd |
| BL1075 | BirA (R118G) | CMV | None | Puro |
| BL1076 | BirA | CMV | None | Puro |
| BL1086 | GFP-MmRab7 | CMV | GFP | Puro |
| BL1087 | Syvn1-2xHA | CMV | HA | Puro |
| BL1099 | ER-GFP (lysozyme-GFP-KDEL) | CMV | GFP | Puro |
| BL1117 | Arih2β-GFP | EF1α | Flag | Bsd |
| BL1118 | GFP-Arih2β | EF1α | Flag | Bsd |
| BL1137 | Arih2 knock in donor template | N.A. | Flag-Avi | Puro |
| BL1139 | Arih2 knockout gRNA 4 | EF1α; U6 | Flag | Puro |
| BL1155 | Arih2β-6xMyc (421aa) | CMV | Myc | Puro |
| BL1156 | Arih2β<sup>C238A</sup>-6xMyc (421aa) | CMV | Myc | Puro |
| BL1192 | Arih2β Δ21-66 | EF1α | Flag | Bsd |
| BL1194 | Arih2β Δ48-66 | EF1α | Flag | Bsd |
| BL1195 | Arih2β Δ41-66 | EF1α | Flag | Bsd |
| BL1196 | Arih2β Δ30-66 | EF1α | Flag | Bsd |
| BL1203 | Smo-3xFlag-Avi | CMV<sup>weak</sup> | Flag-Avi | Bsd |
| BL1206 | Arih1-6xMyc | CMV | Myc | Puro |
| BL1207 | Arih2β Δ22-66 | EF1α | Flag | Bsd |
| BL1208 | Arih2β Δ23-66 | EF1α | Flag | Bsd |
| BL1209 | Arih2β Δ24-66 | EF1α | Flag | Bsd |
| BL1210 | Arih2β Δ25-66 | EF1α | Flag | Bsd |
| BL1211 | Arih2β Δ26-66 | EF1α | Flag | Bsd |
| BL1212 | Arih2β Δ27-66 | EF1α | Flag | Bsd |
| BL1213 | Arih2β Δ28-66 | EF1α | Flag | Bsd |
| BL1214 | Arih2β Δ29-66 | EF1α | Flag | Bsd |
| BL1217 | Arih2β ΔN22 | EF1α | Flag | Bsd |
| BL1218 | N22-GFP-Arih2β | EF1α | Flag | Bsd |
| BL1219 | Arih2βN22-GFP | EF1α | Flag | Bsd |
| BL1231 | AG-PB1 | EF1α | Flag | Bsd |
| BL1232 | Smo-PB1 | CMV | Flag | Bsd |
| BL1233 | Arih2β-3xAG | EF1α | Flag | Bsd |
| BL1234 | Arih2α-AG | EF1α | Flag | Bsd |
| BL1239 | AG | EF1α | Flag | Bsd |
| BL1253 | SmoΔCT-PB1 | CMV | Flag | Bsd |
| BL1254 | Arih2β ΔAriadne-AG | EF1α | Flag | Bsd |
| BL1255 | Arih2β ΔTRIAD-AG | EF1α | Flag | Bsd |
| BL1257 | G-CEPIA1-ER | EF1α | Flag | Bsd |
| BL1258 | G-CEPIA1 | EF1α | Flag | Bsd |
| BL1261 | Arih2β-G-CEPIA | EF1α | Flag | Bsd |
| BL1262 | G-CEPIA-Arih2β | EF1α | Flag | Bsd |
| BL1263 | Smo-3xFlag-GFP-IRES2-mCherry | CMV<sup>weak</sup> | Flag, GFP | None |
| GP778 | 8xGliBox-GgCryD1-nucGFP | Gli | NLS | Nat |
| GP826 | Smo<sup>loop3KRR</sup>-3xFlag | CMV | Flag | Bsd |
| OC166 | lentiCRISPR v2 Nat | EF1α; U6 | Flag | Nat |
| OC173 | Arih2 knockout gRNA 1 | EF1α; U6 | Flag | Puro |
| OC174 | Arih2 knockout gRNA 2 | EF1α; U6 | Flag | Puro |
| PD22 | Smo-3xFlag | CMV | Flag | Bsd |

**Table S2.**
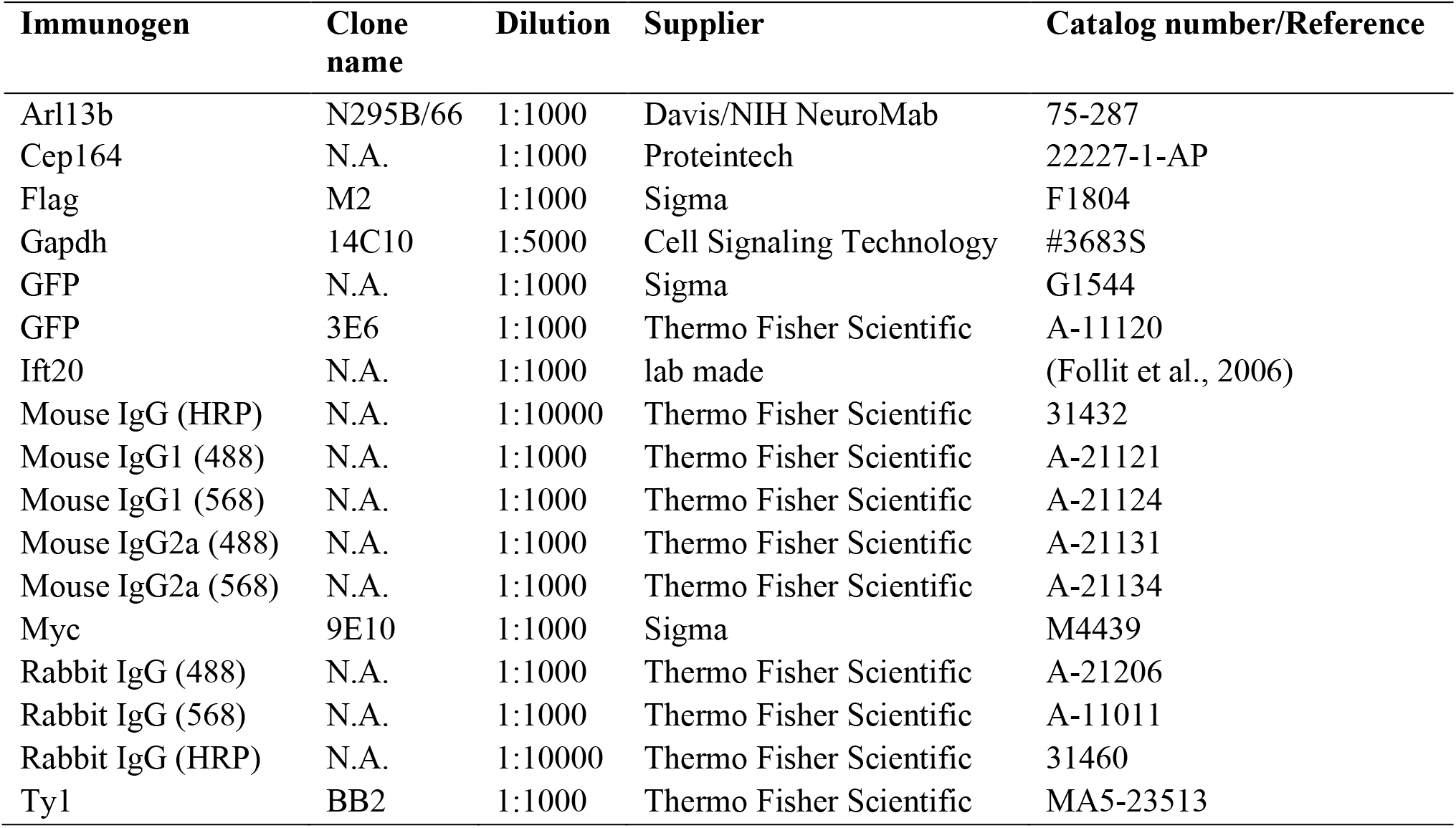
Antibodies.

**Table S3.**
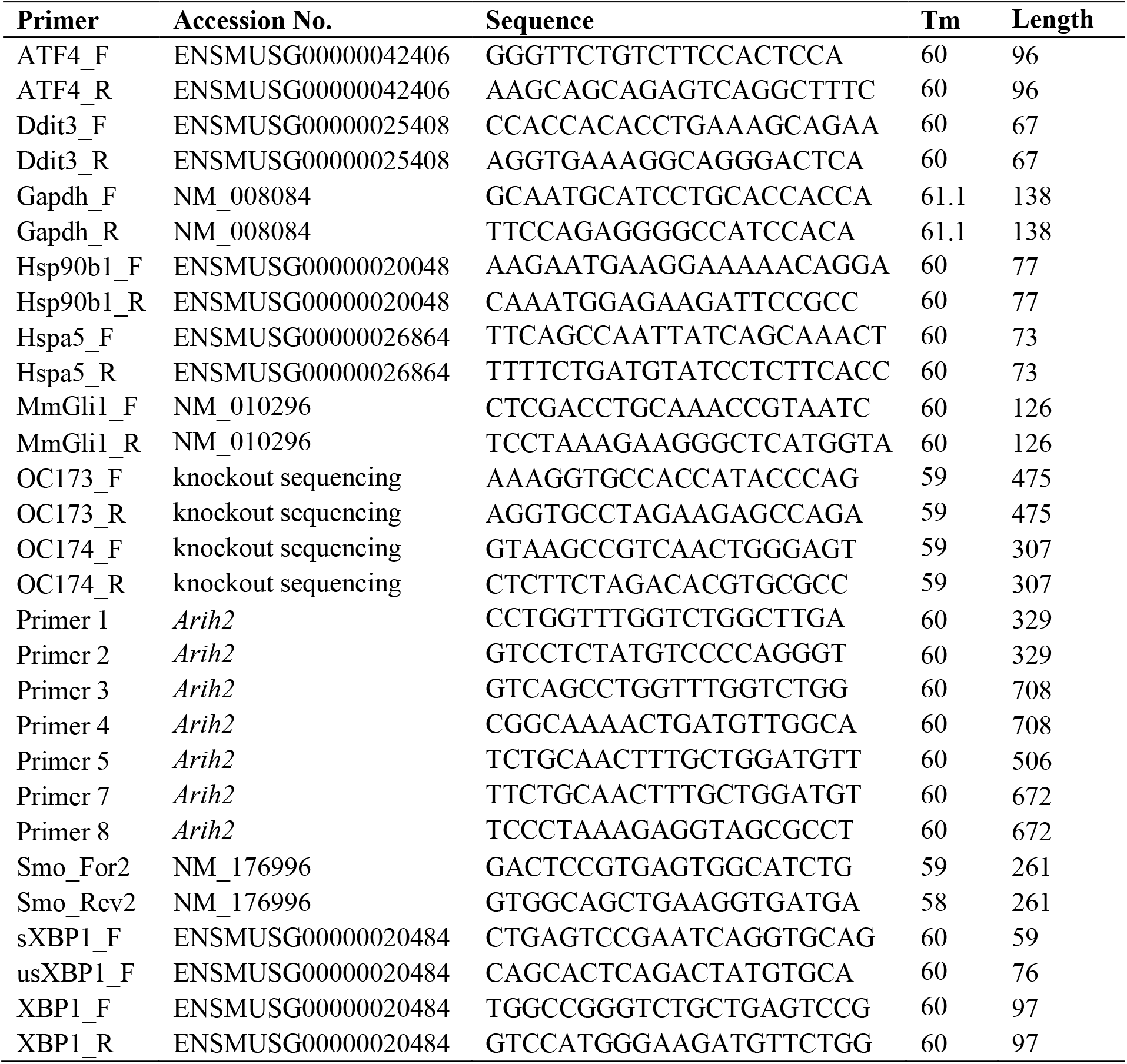
PCR, RT-PCR and Quantitative RT-PCR primers.

